# Thermodynamic analysis of point mutations inhibiting high-temperature reversible oligomer of PDZ3

**DOI:** 10.1101/2020.06.24.169607

**Authors:** T. Saotome, T. Mezaki, B. Subbaian, S. Unzai, J. C. Martinez, S-i. Kidokoro, Y. Kuroda

## Abstract

Differential scanning calorimetry (DSC) indicated that PDZ3 undergoes a peculiar thermal denaturation exhibiting two endothermic peaks due to the formation of reversible oligomers at high temperature (N↔I_6_↔D). This contrasts sharply with the standard 2-state denaturation model observed for small, globular proteins. We performed an alanine scanning analysis by individually mutating three hydrophobic residues at the crystallographic oligomeric interface (Phe340, Leu342, Ile389) and one away from the interface (Leu349, as a control). DSC analysis indicated that PDZ3-F340A and PDZ3-L342A exhibited a single endothermic peak. Furthermore, PDZ3-L342A underwent a perfect 2-state denaturation, as evidenced by the single endothermic peak, and confirmed by detailed DSC analysis, including global fitting of data measured at different protein concentrations. Reversible oligomerization (RO) at high temperatures by small globular proteins is a rare event. While we designed the mutations based on our previous study showing that a point mutation Val380 to a nonhydrophobic amino acid inhibited RO in DEN4 ED3, the results are nevertheless surprising since high-temperature RO involves proteins in a denatured state, as assessed by circular dichroism. Future studies will determine how and why mutations designed using crystal structures determined at ambient temperatures influence the formation of RO at high temperatures, and whether high-temperature ROs are related to the propensity of proteins to aggregate or precipitate at lower temperatures, which would provide a novel and unique way of controlling protein solubility and aggregation.

**Significance:** Despite being a small globular protein, which normaly undergo a two-state unfolding, the thermal denaturation of PSD95-PDZ3, monitored by DSC, exhibited two endothermic peaks. The second peak resulted from a reversible oligomerization (RO) at high temperatures, which is, on its own, a rare phenomenon. In this study, we show that the substitution of a single hydrophobic residue to an alanine at the interface of the crystallographic tetramers inhibited high-temperature RO, resulting in a single endothermic peak. Future studies are required to determine why mutations designed using crystal structures determined at ambient temperatures can inhibit high-temperature RO, and whether the ROs are precursor of irreversible aggregation, If so, the present observations will provide an entirely new basis for creating aggregation-resistant proteins.

## 1. Introduction

Thermal denaturation of small proteins is highly cooperative and reversible, as demonstrated by a sharp sigmoidal curve observed in early spectroscopic measurements [1-5]. Furthermore, differential Scanning microcalorimetry (DSC), which is a prime method for directly determining thermodynamic parameters, shows a single sharp endothermic peak corresponding to thermal transition [6-9]. Thus, a two-state thermal transition between the thermodynamic native (N) and the denatured (D) state is considered as a biophysical hallmark for a natively folded protein [5-6].

The sharp transition is thought to originate from the densely packed, hydrophobic interior of a protein, as exemplified in the jigsaw puzzle model [10]. In small single-domain proteins, the densely packed interior forms a single hydrophobic core that would unpack in a very cooperative manner resulting in the two-state unfolding process. In contrast, thermal denaturation of multi-domain proteins can be multi-stepped, since each domain has a hydrophobic core that can unfold independently of the rest of the protein. Another noteworthy exception to the two-state unfolding of small proteins that have a single hydrophobic core is the molten globule (MG) state. The MG state is a compact globular state with native-like secondary structures, but a loosely packed protein interior [11-14]. The MG state was first observed as an equilibrium or thermodynamic intermediate state at non-physiological and often extreme pHs [11-13], or when artificial mutations destabilizing the hydrophobic core were introduced into the native sequence [15]. Furthermore, the equilibrium MG was associated with kinetic folding intermediates [16-17], but the relationship between the equilibrium and kinetic MG states would benefit from further analyses [18-20]. Thus, under “normal” or near-physiological conditions, a natively folded, single domain with a single hydrophobic core will undergo a two-state thermal transition, and thus its DSC thermogram will appear as a single endothermic peak [5-6], as mentioned above.

Recently, a few single-domain globular proteins (CheY [21], Cro repressor [22], PSD95-PDZ3 [23-24], and DEN4 ED3 [25]) exhibiting DSC thermograms with two endothermic peaks have been reported. Surprisingly, the thermal denaturation is reversible, and the proteins are monomeric both in the native and denatured states, indicating that aggregation was not the reason for the unexpected observation nor for the MG state, which usually appears only under extreme conditions. The origin of the unexpected thermal transition was the formation of a reversible oligomer at high temperature that we named the “reversible oligomerization” (RO) state. DSC thermograms showed two endothermic peaks, where the first peak was associated with the unfolding of the protein and its oligomerization (N↔I_n_), and the second peak reflected the dissociation of the intermediate oligomer into a monomeric denatured state (I_n_↔D).

A previous study using DEN4 ED3 suggested that the hydrophobic interaction at the oligomeric interface in the crystal structure of DEN4-ED3 was responsible for RO formation at high temperatures [25]. DEN4 ED3 is the third domain of an envelope protein derived from Dengue virus type 4, with a molecular weight of 11.4 kDa. It has an immunoglobulin fold composed of 9 β-strands. The immunoglobulin-fold or β-sandwich fold is one of the most common folds, and its thermal denaturation usually follows a two-state model [26], but the DSC thermogram of DEN4 ED3 exhibited two endothermic peaks and indicated the presence of a tetrameric intermediate state (N↔I_4_↔D). We showed that the second peak disappeared when Val380, which is located at the crystallographic oligomer interface, was replaced with a non-hydrophobic residue (Ala, Ser, Thr, Asn, or Lys). This indicated that the formation of the high-temperature RO by DEN4 ED3 was, surprisingly, inhibited by mutating a single hydrophobic residue at the interface of its crystallographic structure, which was solved at low temperatures.

PDZ3 is the third PDZ domain of a synaptic carrier protein, PSD95, with a molecular weight of 11.0 kDa. It is made of three α-helices and eight β-strands. DSC thermograms of PDZ3 also show two endothermic peaks, which suggested the formation of a high-temperature RO state (N↔I_n_↔D) [23-24]. Furthermore, earlier work on PDZ3 indicated that the ROs might be precursors of heat aggregation. Hence, strategies for inhibiting formation of RO might facilitate the design of proteins resistant to aggregation and precipitation.

In the present study, we attempted to inhibit the formation of high-temperature RO in PDZ3 by introducing a single mutation at the oligomeric interface of PSD95-PDZ3 (PDZ3) as we succeeded in doing with DEN4 ED3. To this end, we analyzed the X-ray crystal-structure of PDZ3, which formed a tetramer in the unit cell [27]. The Buried Surface Area in the tetrameric crystal structure of PDZ3 (PDB ID: 3I4W) suggested that Phe340/Leu342/Ile389 participated in oligomerization of protein monomers within the crystal structure. Hence, we replaced four hydrophobic amino acids (Phe340, Leu342, Ile389, Leu349) with alanine and designed four single-mutated PDZ3 variants. Eventually, L342A yielded a mutant that denatured according to a two-state model. This finding shows that it is possible to avoid formation of RO using single mutations that do not disturb the native state structure. It also validated our hypothesis that the RO state can be abolished by a single mutation of a hydrophobic residue at the oligomeric interface of the crystal structure.

## 2. Materials and Methods

### 2-1. DSSP and PDBePISA analysis

Accessible Surface Area (ASA) was calculated using X-ray crystallographic data of PSD95-PDZ3 (PDB ID: 3I4W) by DSSP [28]. The four monomeric chains in each asymmetric unit cell form a crystallographic tetramer. Buried Surface Area (BSA) was determined by calculating the ASA of the tetrameric unit and subtracting the ASA of the monomeric structure.

Additionally, X-ray crystallographic data of PSD95-PDZ3 were also analyzed using PDBePISA (https://www.ebi.ac.uk/pdbe/pisa/) [29]. BSA values differed between DSSP and PDBePISA, because DSSP analyzes the tetramer’s interface, whereas PDBePISA analyzes the interface of dimers and calculates three parameters (ASA, BSA, Δ^i^*G*,). Δ^i^*G* (solvation free energy) calculated by PDBePISA indicates the solvation energy of the corresponding residue in kcal/M between the dissociated and the associated structures. Hydrogen bonds, salt bridges, and disulfide bonds are included in calculation ofΔ^i^*G* (−0.44, −0.15 and −4 kcal/mol per bond, respectively).

### 2-2. Synthesis, expression, and purification

Synthetic genes encoding PDZ3 were cloned into a pBAT4 vector, and single mutations were introduced by site-directed mutagenesis using a QuikChange protocol (Stratagene, USA). All variants were overexpressed in *Escherichia coli* strain BL21 (DE3) with 1 L of LB medium. Protein expression was induced by addition of 0.2 mM IPTG when the OD reached 0.6. After centrifugation, harvested cells were lysed in 50 mM Tris-HCl (pH 8.7) by ultrasonication. Next, the supernatant fraction of the cell lysate was acidified to pH 3 by adding several drops of 1 M HCl. Finally, proteins were purified by reverse-phase HPLC, lyophilized, and stored at −30°C until further use.

### 2-3. Mass Spectrometry (MALDI-TOF MS) Measurements

Matrix-Assisted Laser Desorption/Ionization-Time of Flight (MALDI-TOF) MS measurements were performed by using the plate with autoflex speed TOF/TOF (Bruker Daltonics). The matrix solution was prepared by dissolving 10 mg of sinapic acid in 1 mL of a solution containing 300 μL of acetonitrile, 100 μL of 1% trifluoroacetic acid and 600 μL of MQ. Samples for MALDI-TOF MS were prepared by mixing 1 μL of protein solution with 4 μL of the matrix solution. 1 μL of sample mixtures were spotted and air-dried on the MALDI-TOF MS plate.

### 2-4. Differential Scanning Calorimetry (DSC) measurements

Samples were prepared by dissolving lyophilized proteins in Milli Q water (MQ) and dialyzed for 18 hours at 4°C in 50 mM potassium-phosphate buffer (pH 7.5) using a Spectra/Por 3 membrane (MWCO of 3.5 kDa) with one buffer exchange. After dialysis, the protein concentrations of samples were adjusted to 1 mg/mL, and samples were filtered with 0.20 μm membrane filters (Millex-GV; Millipore, USA) to remove aggregates, followed by degassing of the samples. Protein concentrations and pHs of the samples were confirmed just before performing experiments.

DSC measurements were performed using a VP-DSC MicroCalorimeter (MicroCal, USA) at a scan rate of 1.0°C / min in the temperature range of 20 to 100°C. Blank measurements were taken using 50 mM potassium-phosphate buffer (pH 7.5) several times before measurements. The reversibility of the thermal unfolding of the protein was checked by repeated scans of the same sample. Thermodynamic parameters (*T*_m_ and Δ*H* (*T*_m_)) were determined by analyzing the apparent heat capacity curves using a non-linear, least-square fitting algorithm, DDCL3 [30-31], and assuming a linear temperature dependence of the heat capacity for the native and denatured states.

### 2-5. Circular Dichroism (CD) measurements

Samples were prepared by dissolving lyophilized proteins in MQ, and protein concentrations of samples was adjusted to 0.5 mg/mL in 50 mM potassium-phosphate buffer (pH 7.5). Samples were filtered with 0.20 μm membrane filters (Millex-GV; Millipore, USA) to remove aggregates. Protein concentrations and pHs of samples were confirmed just before performing experiments.

CD measurements were conducted using a JASCO J-820 spectropolarimeter at 0.2 mg/mL protein and 50 mM potassium-phosphate buffer (pH 7.5) using quartz cuvettes with a 2-mm optical path length. CD spectra were obtained by using the CD value in the range of 200-260 nm. Thermal stability was measured at a protein concentration of 0.5 mg/mL in 50 mM potassium-phosphate buffer (pH 7.5), at a 1°C/min scan rate, and monitored between 20°C and 90°C using the CD value at 220 nm. Midpoint temperatures (*T*_m_) were computed by means of least-squares fittings of experimental data with a two-state model using Origin 6.1 J.

### 2-6. Dynamic Light Scattering (DLS) measurements

Samples were prepared by dissolving lyophilized proteins in MQ, and protein concentrations of samples were adjusted to 1 mg/mL in 50 mM potassium-phosphate buffer (pH 7.5). Samples were filtered with 0.20 μm membrane filters (Millex-GV; Millipore, USA) to remove aggregates. Protein concentrations and pHs of samples were confirmed just before performing experiments.

DLS measurements were performed by using a glass cuvette with a Zeta-nanosizer (Nano S; Malvern). Sample temperatures were increased from 25°C to 60°C, and then to 90°C, before being cooled to 25°C to assess reversibility. The hydrodynamic radius (*R*_h_) was calculated from size-volume graphs using the Stokes-Einstein equation.

### 2-7. Analytical Ultracentrifugation (AUC) measurements

Samples were prepared by dissolving lyophilized proteins in MQ, and protein concentrations of samples were adjusted to 1 mg/mL in 50 mM potassium-phosphate buffer (pH 7.5). Samples were filtered with 0.20 μm membrane filters (Millex-GV; Millipore, USA) to remove aggregates. Protein concentrations and pHs of samples were confirmed just before performing experiments.

Sedimentation velocity experiments were carried out using an Optima XL-A analytical ultracentrifuge (Beckman–Coulter) with a four-hole An60Ti rotor at 25°C. Before centrifugation, samples were dialyzed overnight against 50 mM potassium-phosphate buffer (pH 7.5). Each sample was then transferred into a 12-mm double-sector Epon cell and centrifuged at a rotor speed of 50,000 rpm. Concentrations were monitored at 280 nm. Sedimentation velocity data were analyzed using the continuous distribution c(s) analysis module in the SEDFIT program [32]. The range of sedimentation coefficients where the main peak was present was integrated to obtain the weighted average sedimentation coefficient. The c(s) distribution was converted into c(M), a molar mass distribution. Solvent density, viscosity, and protein partial specific volumes were calculated using SEDNTERP software [33].

## 3. Results

### 3-1. Two endothermic peaks in the wt-PDZ3 thermogram

Although thermal denaturation monitored by CD exhibited a sigmoidal curve typical of a natively folded protein, the DSC thermogram of PDZ3-wt exhibited two endothermic peaks, as reported previously [23-24] (Figure 3). The apparent melting temperature, defined by the maximum of the first endothermic peak, decreased when the protein concentration was increased (Figure 3, Supplementary Table S5). This dependency of the melting temperature on protein concentration is contrary to that observed for proteins that form oligomers in the native state [34-35] and imply the presence of an oligomeric intermediate state at high temperature. The second endothermic peak is attributed to the dissociation of the high-temperature oligomeric state into a monomeric, unfolded state.

**Figure 1:**
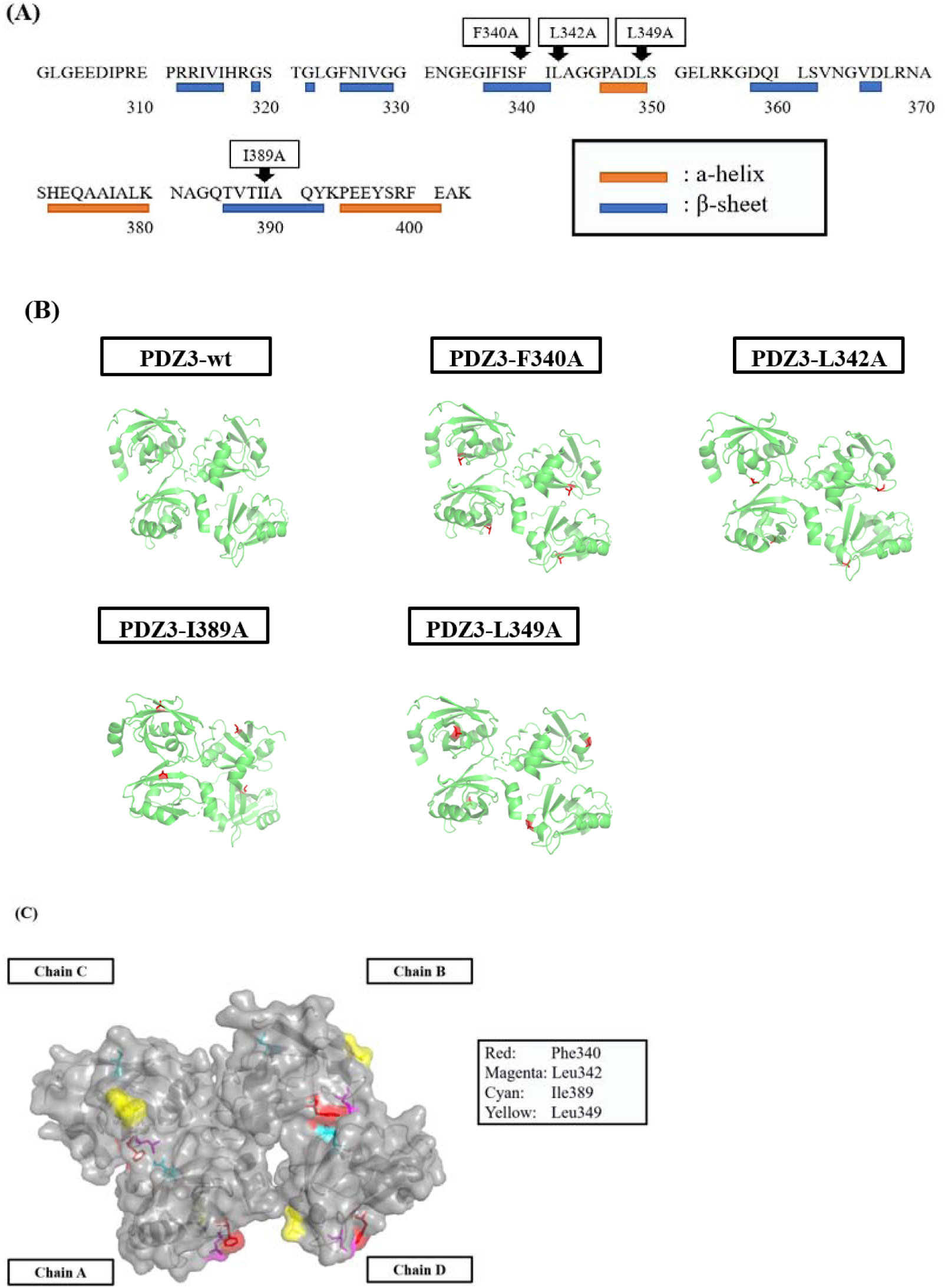
Amino acid sequence (A) and ribbon model (B) of PDZ3 variants. The ribbon model was drawn using Pymol and PDB structure 3I4W. The sites of single alanine mutations are shown in red. (C) The residual conformation of the interface of PDZ3-wt (PDB ID: 3I4W). Two polypeptide chains are represented as wireframe models using Pymol. The backbone of PDZ3 (Right grey), Phe340 (Red), Leu342 (Magenta), Ile389 (Cyan) and Leu349 (Yellow) are distinguished by different colors.

**Figure 2:**
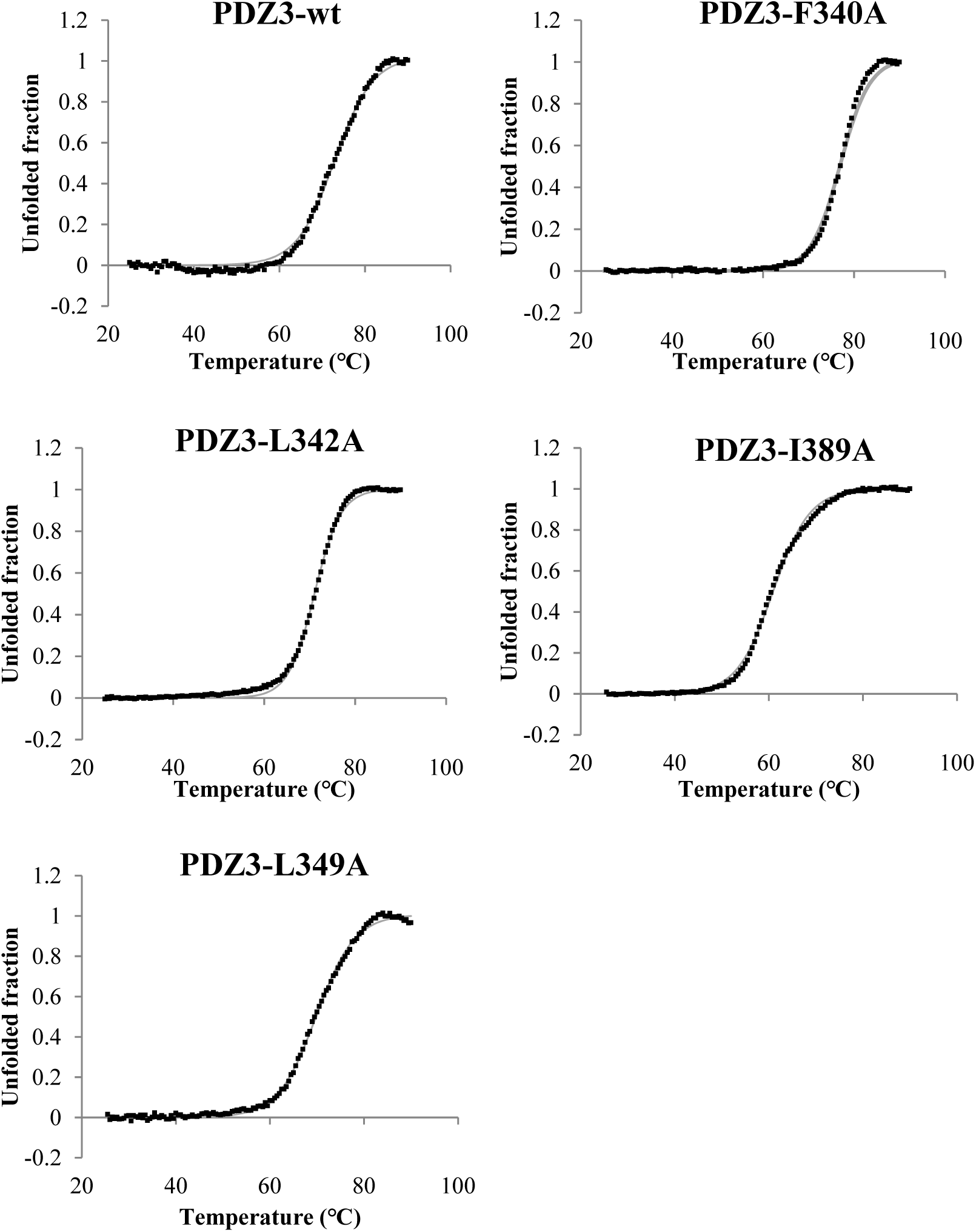
CD thermal denaturation curves of PDZ3 variants at 0.5 mg/mL and pH 7.5. Samples were heated at the scan rate of +1°C/min, and CD values were monitored at 220 nm. Black dots and gray lines represent the raw CD value and the fitting curves.

**Figure 3:**
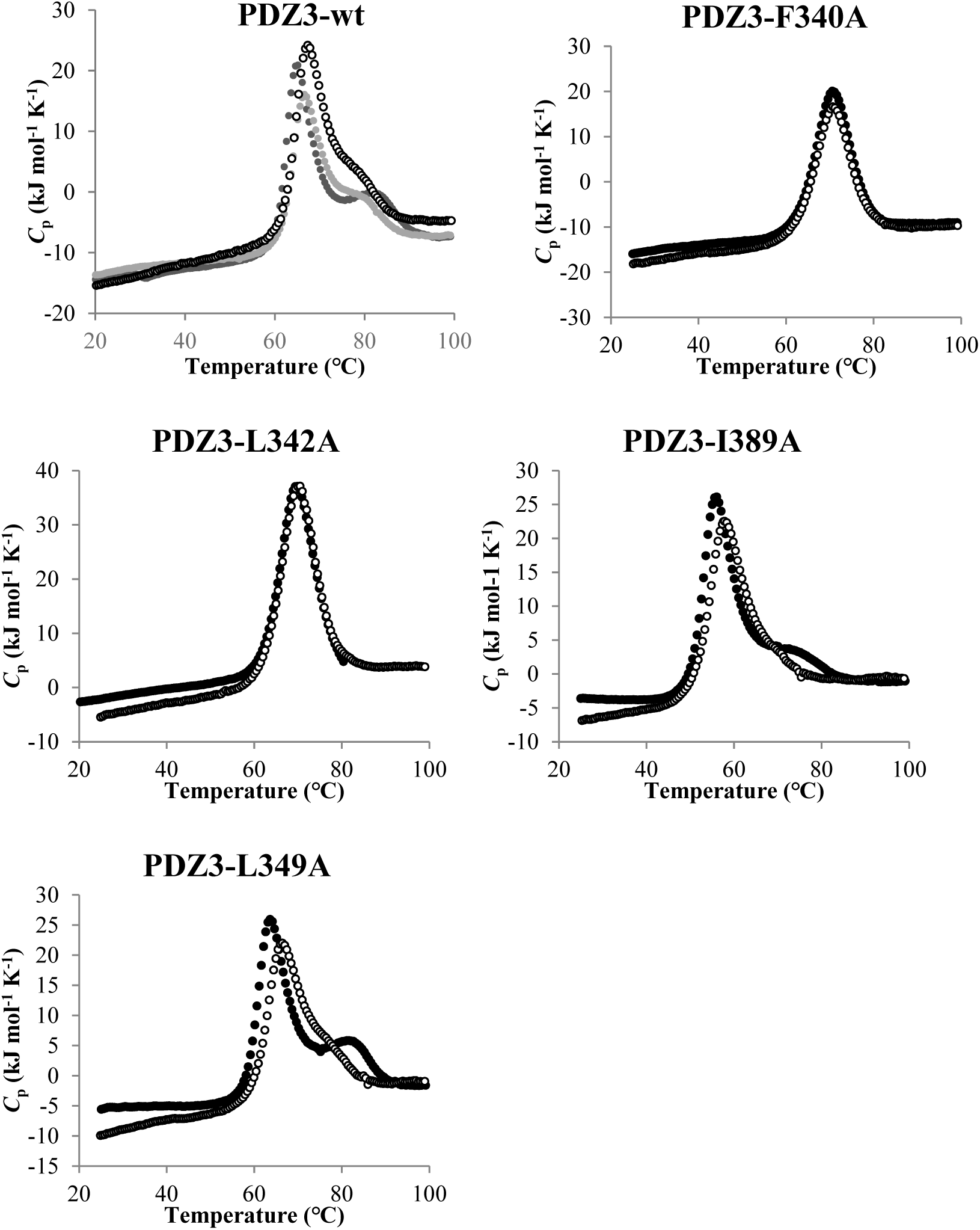
Concentration dependence of DSC thermograms of PDZ3 variants at 0.5-1 mg/mL, pH 7.5 and 1°C /min scan rate. Open black circle, open gray circle and open white circle show DSC thermograms at protein concentrations of 1 mg/mL, 0.75 mg/mL and 0.5 mg/mL.

### 3-2. Design of PDZ3 variants

We designed point mutations in PDZ3 intended to inhibit formation of high-temperature reversible oligomers. Consistent with our design strategy developed using DEN4 ED3, we searched for hydrophobic residues at the crystal interface and replaced them with alanine. To this end, we used the crystal structure of the wild type PDZ (PDZ3-wt), which contains a PDZ tetramer in the unit cell (Fig 1, Table 1, Supplementary Table S1). Based on our previous analysis using DEN4-ED3, we assumed that high-temperature oligomers were generated by hydrophobic patches on the protein that would interact with the nearby protein. Hence, we focused on hydrophobic residues that exhibited the largest BSA, according to DSSP. BSA was determined by subtracting the ASA of the tetrameric unit from the ASA of the monomeric structure. Here we took Ile, Leu, Val, Phe as hydrophobic, as defined by the Kyte and Doolittle hydropathy scale [36], and we selected three hydrophobic residues, Phe 340, Leu342, and Ile389, exhibiting the largest BSA (Table 1, Supplementary Table S1).

**Table 1.** Accessible and buried surface areas of hydrophobic residues of PDZ3-wt. Residues with ASA> 0 Å^2^ of at least one of the chains are listed. Hydrophobic residues (Ile, Leu, Val, Phe) were selected using the hydropathy scale of Kyte and Doolittle [36]. BSA (Buried Surface Area) values of the four polypeptide chains (A, B, C, D) of PDZ3-wt (PDB ID: 3I4W) at the crystallographic tetramer calculated by DSSP. Values of BSA were obtained by subtracting the ASA (Accessible Surface Area) in the tetrameric structure from the ASA calculated from the monomer. The largest ASA and BSA are highlighted in brown, and selected residues are underlined.

| Residue | ASA ($\text{\AA}^2$) of monomer | | | | BSA ($\text{\AA}^2$) of tetramer | | | |
| --- | --- | --- | --- | --- | --- | --- | --- | --- |
|  | A | B | C | D | A | B | C | D |
| ILE 307 | 32 | 39 | 22 | 40 | 25 | 0 | 0 | 0 |
| ILE 314 | 1 | 1 | 1 | 3 | 0 | 0 | 0 | 0 |
| VAL 315 | 50 | 52 | 50 | 53 | 0 | 0 | 0 | 3 |
| ILE 316 | 3 | 3 | 3 | 4 | 0 | 0 | 0 | 0 |
| LEU 323 | 13 | 13 | 12 | 14 | 0 | 5 | 0 | 0 |
| PHE 325 | 19 | 21 | 20 | 21 | 0 | 0 | 0 | 0 |
| ILE 327 | 21 | 23 | 19 | 23 | 0 | 2 | 7 | 0 |
| VAL 328 | 34 | 35 | 33 | 33 | 0 | 33 | 31 | 0 |
| <u>PHE 340</u> | 98 | 98 | 99 | 97 | 0 | 64 | 63 | 0 |
| ILE 341 | 41 | 35 | 41 | 38 | 0 | 0 | 0 | 0 |
| <u>LEU 342</u> | 60 | 63 | 59 | 61 | 0 | 59 | 55 | 0 |
| <u>LEU 349</u> | 143 | 127 | 136 | 127 | 0 | 0 | 0 | 0 |
| LEU 353 | 11 | 11 | 11 | 11 | 0 | 0 | 0 | 0 |
| LEU 360 | 23 | 19 | 20 | 20 | 20 | 0 | 0 | 18 |
| VAL 362 | 1 | 1 | 1 | 1 | 0 | 0 | 0 | 0 |
| VAL 365 | 74 | 75 | 75 | 76 | 0 | 0 | 0 | 0 |
| LEU 367 | 3 | 3 | 4 | 3 | 0 | 0 | 0 | 0 |
| ILE 377 | 73 | 75 | 71 | 73 | 0 | 12 | 0 | 0 |
| LEU 379 | 16 | 14 | 14 | 14 | 0 | 2 | 0 | 0 |
| <u>ILE 389</u> | 50 | 48 | 49 | 48 | 46 | 0 | 0 | 45 |
| PHE 400 | 90 | 92 | 89 | 89 | 0 | 19 | 0 | 0 |

The above design was confirmed using PDBePISA (Supplementary Fig S1, Supplementary Table S2). Candidate residues were identified at the two largest interfaces “ID-1 (average 518.0 Å^2^)” and “ID-2 (average 449.6 Å^2^)” (Supplementary Table S3) predicted by PDBePISA. According to PDBePISA, Ser371, Glu373, and Glu331, and Leu342, Ile 389, and Phe 340 are critical for maintaining interfaces ID1 and ID2. Phe 340, Leu342, and Ile 389, which were the same residues selected using DSSP. Additionally, we chose, as a control, a hydrophobic residue (Leu 349) with a large ASA but with a negligible BSA, meaning that this residue is not buried by formation of the tetramer (Supplementary Table S2).

### 3-3. Unaltered physicochemical properties of PDZ3 variants

The single alanine substitution on PDZ3 did not change the native structure nor its physicochemical properties at low temperature up to ∼50°C. In particular, all PDZ3 variants were monomers at 25°C, as shown by AUC (Supplementary Fig S2, Table 2), and CD spectra of all variants were nearly identical at 25°C. (Supplementary Fig S3). Additionally, the thermal denaturation curve developed by monitoring CD at 220 nm showed a typical sigmoidal transition that could be fitted by a two-state model (N↔D) (Fig 2, Table 3). However, PDZ3-I389A was 10°C less stable than the other variants, indicating that a single mutation could affect the thermal stability of PDZ3, as in our previous observations with BPTI [37].

**Table 2.**
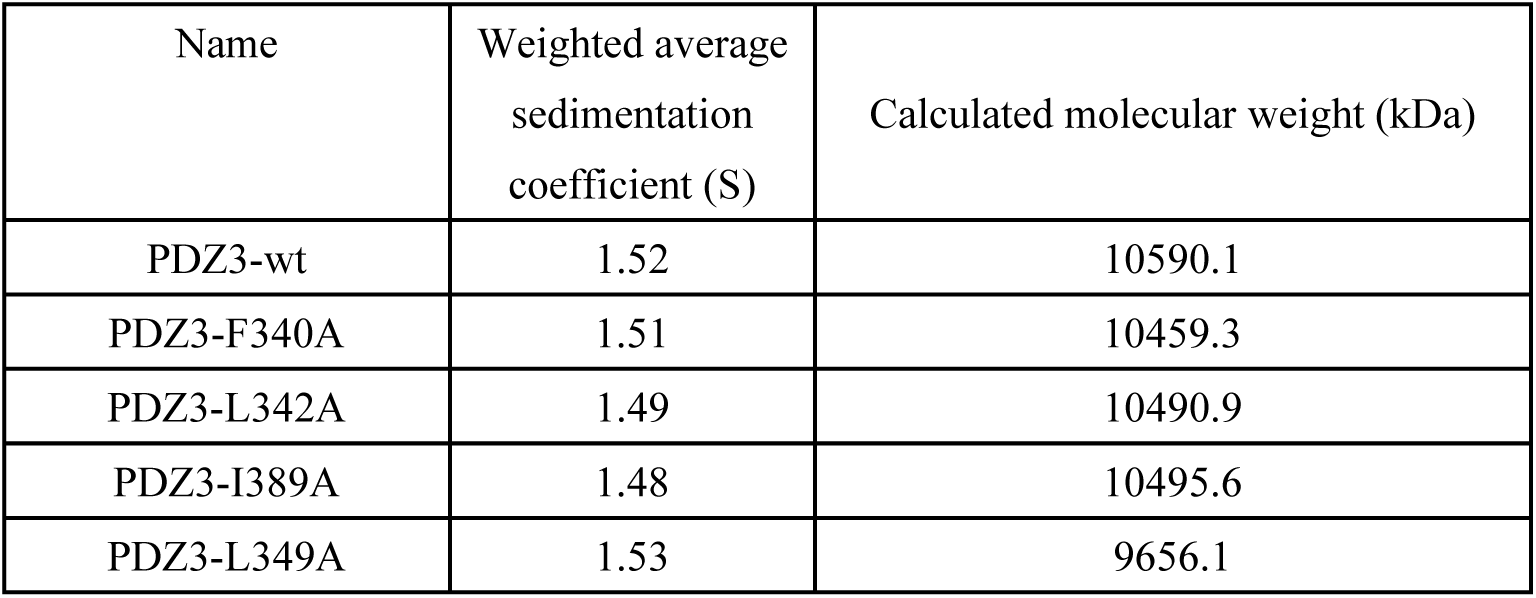
The sedimentation coefficient and calculated molecular weight of PDZ3 variants in AUC measurements at 1 mg/mL, pH 7.5 and 25°C. Sedimentation velocity analysis of PDZ3 variants was performed at 25°C and experimental data were analyzed using SEDFIT [32] and SEDNTERP [33].

**Table 3.** Midpoint temperature (*T*_m_) and van’t Hoff enthalpy (Δ*H*_van’t Hoff_ (*T*_m_)) of PDZ3 variants by two-state analysis in CD measurement at 0.5 mg/mL, pH 7.5 and 1°C/min scan rate. These thermodynamic parameters were calculated by fitting thermal denaturation curves of PDZ3 variants.

| Name | $T_m$ (°C) | $\Delta H_{\text{van't Hoff}}(T_m)$ (kJ/mol) |
| --- | --- | --- |
| PDZ3-wt | 72.58 | 249.13 |
| PDZ3-F340A | 76.68 | 392.04 |
| PDZ3-L342A | 70.97 | 339.13 |
| PDZ3-I389A | 60.99 | 237.23 |
| PDZ3-L349A | 69.74 | 241.75 |

### 3-4. Inhibition of high-temperature RO by a single mutation

DSC indicated that our design for abolishing the RO state was successful. In particular, PDZ3-F340A and PDZ3-L342A at a 1 mg/ml concentration and pH 7.5 showed a single endothermic peak, and PDZ3-I389A showed a large peak overlapped with a small shoulder (Fig 3, Supplementary Table S5, Supplementary Table S5). The mutations changed the molar fraction of intermediate oligomer (Fig 4, Supplementary Fig S5), as well as other thermodynamic properties related to the formation of the RO (Supplementary Table S5). On the other hand, PDZ3-L349A, which is a surface-exposed hydrophobic residue located far from the crystal oligomeric interface, showed two endothermic peaks almost identical to PDZ3-wt.

**Figure 4.**
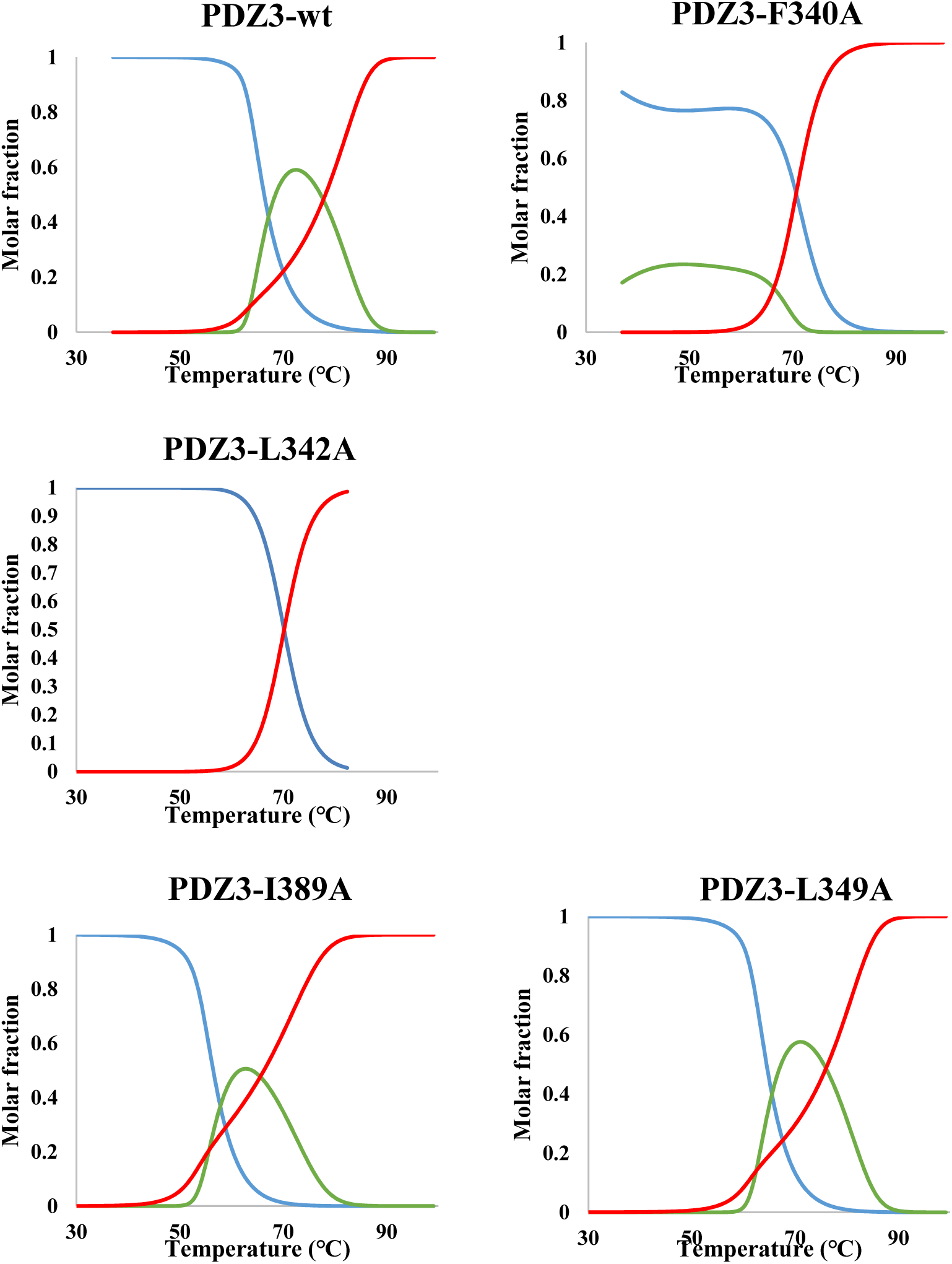
Molar fraction of PDZ3-wt (N-I_6_-D model), PDZ3-F340A (N-I_4_-D model), PDZ3-L342A (N-D model), PDZ3-I389A (N-I_5_-D model), and PDZ3-L349A (N-I_5_-D model) by DDCL3 analysis of DSC thermograms at 1 mg/mL, pH 7.5 and 1°C /min scan rate. Natively folded monomer (Blue); Intermediate oligomer (Green); Unfolded monomer (Red).

The DSC results were corroborated by hydrodynamic radii measured by DLS size-volume analysis. DLS indicated that RO formation at high temperature was significantly inhibited for PDZ3-F340A and PDZ3-L342A (Supplementary Fig S6, Table 5), which showed single endothermic peaks. Their hydrodynamic radii, *R*_h_, were almost unchanged by heating from 25°C to 60°C (1.76-1.77 nm), in contrast to the *R*_h_ of PDZ3-wt, PDZ3-L349A and PDZ3-I389A, which increased at 60°C to 2.40 nm, 4.07 nm and 2.20 nm respectively (Table 5). Altogether, these observations indicate that high-temperature RO is inhibited or significantly reduced by alanine substitutions of single superficial hydrophobic residues located at the crystallographic oligomer interface.

**Table 4.**
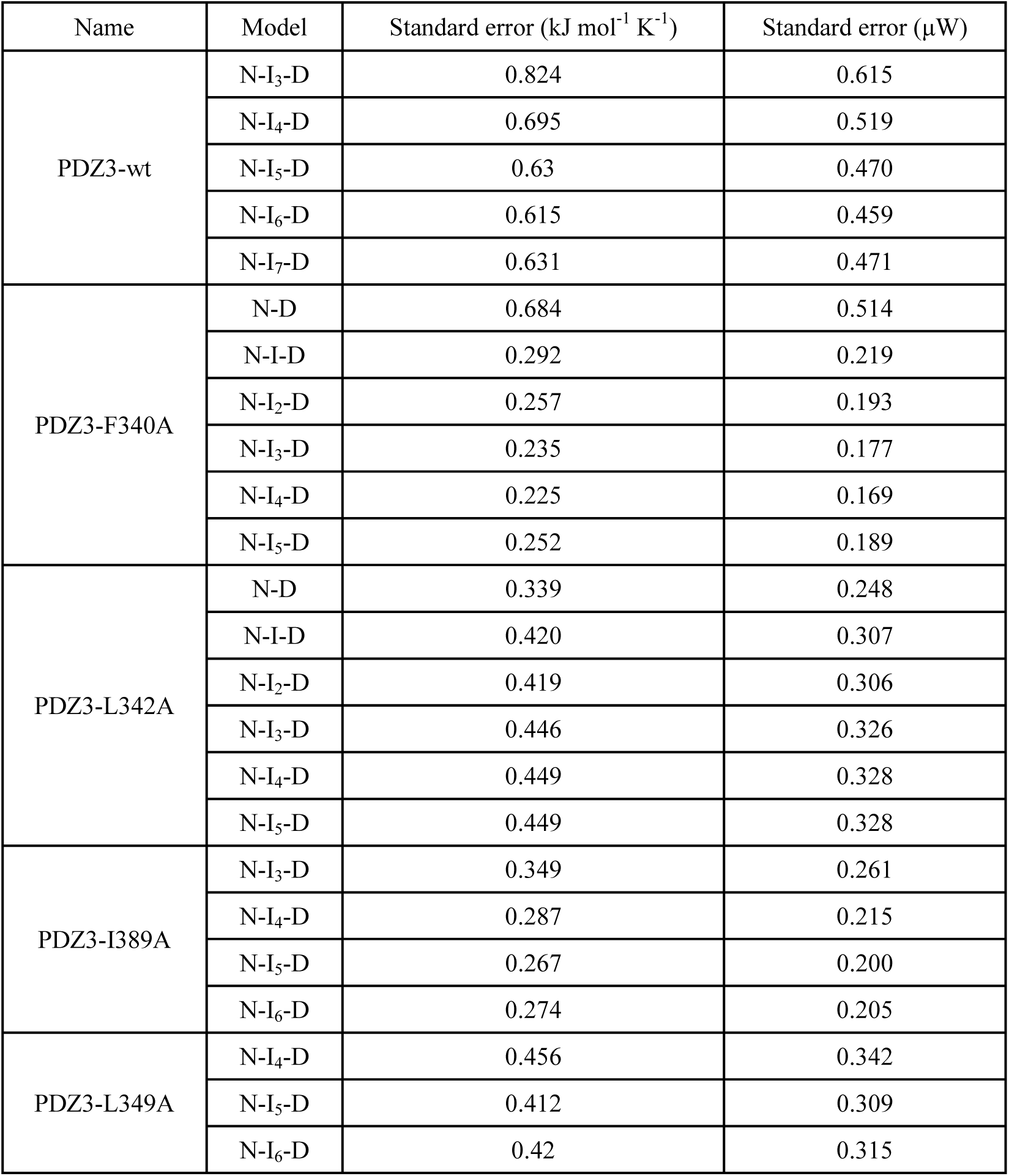
Standard errors of PDZ3 variants between DSC raw data and fitting curves calculated with DDCL3. The oligomer number of the intermediate (I) state varied from monomer (I) to heptamer (I_7_) at the thermal transition models. Optimal thermal transition models of each PDZ3 variant were determined by comparing the minimum standard error.

**Table 5.**
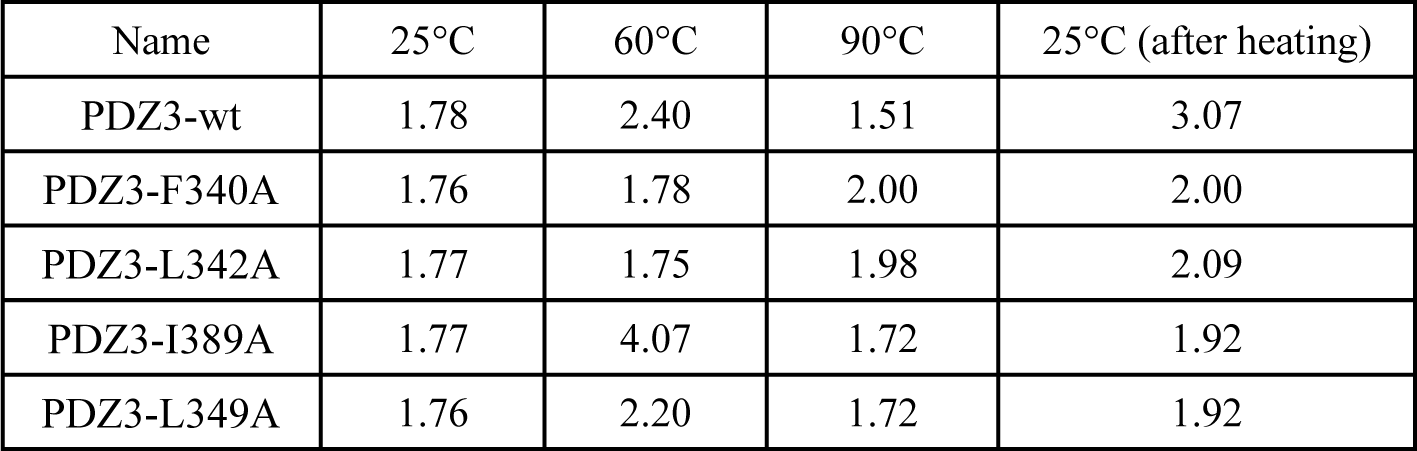
Hydrodynamic radius (nm, *R*_h_) of PDZ3 variants by DLS measurements at 1 mg/mL, pH 7.5 and 25-90°C. *R*_h_ values were calculated from size-volume graphs at 25°C, 60°C, 90°C and 25°C after heating.

### 3-5. Thermodynamic parameters of PDZ3 and its variants

We performed a global fitting of the DSC thermograms measured at multiple protein concentrations in order to provide a detailed characterization of the thermal transition of PDZ3 variants. Both PDZ3-F340A and PDZ3-L342A showed a single endothermic peak. However, global fitting indicated that only the latter followed a perfect two-state denaturation model (N↔D): The ratio of the calorimetric and van’t Hoff enthalpy of PDZ3-L342A was essentially 1.0, indicating the absence of an intermediate state (Table 3, Supplementary Table S5). Additionally, the residuals of global fittings were almost identical between the two-state (N↔D) and three-state models (N↔I_n_↔D). Hence, the thermal transition of PDZ3-L342A was perfectly described by a two-state model (Table 4); thus, the formation of the high-temperature RO was fully inhibited.

Though the DSC thermogram of PDZ3-F340A, showed a single endothermic peak, the global fitting analysis indicated a three-state (N↔I_n_↔D) denaturation. Indeed, the standard error of the global fitting drastically decreased using a three-state model, but the molar fraction of the intermediate state (I_n_) was very small. Thus, F340A strongly, but not fully inhibited formation of the high-temperature RO (Fig 4, Supplementary Fig S5, Supplementary Table S5).

PDZ3-I389A and PDZ3-L342A obviously could not be fitted using a two-state model, because two endothermic peaks appeared in their DSC thermograms. The second endothermic peak of PDZ3-I389A was slightly smaller than PDZ3-wt, whereas that of PDZ3-L349A was almost identical with PDZ3-wt (Supplementary Fig S5, Supplementary Table S5). Thus, I389A and L349A did not significantly inhibit formation of high-temperature ROs.

## Discussion

PSD95-PDZ3’s DSC thermogram exhibits two endothermic peaks, which is atypical of the denaturation curve observed in single-domain, monomeric, globular proteins. Such an observation is rare, but it was reported for another small, single-domain protein, DEN4 ED3, where the second endothermic peak resulted from high-temperature reversible oligomerization. Here, our DSC analysis showed that PDZ3 undergoes thermal denaturation with a hexameric, high-temperature, RO intermediate state (N↔I_6_↔D). To date, reversibility is unusual when denatured proteins oligomerize, because this often triggers irreversible aggregation followed by precipitation, exhibiting a strong exothermic peak during DSC measurement. In addition, high-temperature RO has rarely been reported, precisely because it occurs at high temperatures, which we speculate might have made it difficult to observe.

For the purpose of discussion, let us note that it is counter-intuitive that a mutation designed using the crystal structure, specifically the replacement of a hydrophobic residue at a crystallographic interface, would drastically affect the formation of unfolded oligomers at high temperature. However, this can be rationalized by assuming that the high-temperature RO occurs through oligomerization of folded proteins, either in a native state or more likely, in an MG-like state, where the hydrophobic core is conserved but with highly dynamic sidechains [11-20]. Thus, although further studies are required to fully confirm this hypothesis, we speculate that PDZ3-wt oligomerizes in a folded state, where the interactions observed in the crystal structure prevail (either native or MG-like). Unfolding would occur after oligomerization, as confirmed by comparing CD and DSC data (Figs. 2 and 3), which increases the destabilizing effect through reversed hydrophobicity [38] becoming stronger at high temperatures [39], as we showed for oligomerization of tagged BPTIs [Nakamura et al. submitted]. This interpretation is corroborated by our present design of PDZ3-F340A and PDZ3-L342A, where alanine substitution of hydrophobic residues at the crystallographic oligomer interface inhibited RO formation, but not PDZ3-L349A, which was surface exposed, but located far from the oligomerization site. Furthermore, a recent study revealed that the Ebola virus matrix protein VP40 also shows a DSC thermogram with two endothermic peaks. In addition, a single mutation (L117R) in the hydrophobic region of the N-terminal domain at the dimeric crystallographic interface surprisingly suppressed the second endothermic peak [40]. This observation also corroborates our present finding that the hydrophobic interaction at the interface of the natively-folded oligomer actively contributes to formation of high-temperature RO.

Finally, let us note that single mutations in recombinant proteins that drastically alter their physicochemical properties are rare because multiple interdependent factors (compensation of enthalpy and entropy [41-42], hydrogen bonds [43-44], salt bridges [45-46] and disulfide bonds [41, 47]) are usually involved. Mutations often impair the original structure or function, with a few exceptions [48-49]. Thus, a single alanine substitutions on PDZ3 fullyinhibiting the formation of high-temperature is surprising, and may be because interaction energies stabilizing the RO state are weak.

## 4. Conclusion

Although PSD95-PDZ3-wt is a monomeric single-domain protein, it shows a peculiar thermal denaturation curve with two endothermic peaks, as observed by DSC. Such observations are rare, but this is the second example, along with our previous report of DEN4 ED3. Hence, RO formation might be a phenomenon common to many small globular proteins, but was overlooked so far, because it occurs at high temperature.

Finally, our strategy for inhibiting RO using the crystal structure of PDZ3, indicated that a single alanine substitution of a hydrophobic residue, L342A, located at the monomeric surface fully inhibited RO formation at high temperature. On the other hand, a control substitution replacing Leu349, an exposed hydrophobic residue far from the oligomerization site indeed did not inhibit RO formation, indicating that RO formation must be associated with hydrophobic residues at the interface of the crystallographic oligomers. Moreover, RO formation at high temperature may be a precursor of aggregation in the case of PDZ3, and our design strategy may have a practical usefulness in designing aggregation-resistant proteins.

## Abbreviations

PSD95: postsynaptic density-95 protein;
PDZ3: the third PDZ domain from PSD95;
RO: Reversible Oligomer;
DSC: Differential Scanning Calorimetry;
PISA: Proteins Interfaces Surfaces and Assemblies;
AUC: Analytical Ultracentrifugation;
CD: Circular Dichroism;
DLS: Dynamic Light Scattering;
*R*_h_: Hydrodynamic Radius;
*T*_m_: Midpoint Temperature;

## Author contributions

ST, SK, and YK designed the study and wrote the manuscript. ST, TM, and SK carried out the DSC analysis. ST and SB prepared the samples to perform the spectroscopic analysis. JM provided the materials and co-wrote the manuscript, SU carried out the AUC analysis. All authors contributed to the editing of the manuscript and approved the final version.

## Acknowledgments

We thank all members of the Kuroda Laboratory for discussion and technical assistance. We are grateful to Professors Tsuyoshi Tanaka and Tomoko Yoshino for the use of ZetaNanosizer.

